# Re-evaluation of a Tn*5*::*gacA* mutant of *Pseudomonas syringae* pv. *tomato* DC3000 uncovers roles for *uvrC* and *anmK* in promoting virulence

**DOI:** 10.1101/774711

**Authors:** Megan R. O’Malley, Alexandra J. Weisberg, Jeff H. Chang, Jeffrey C. Anderson

**Affiliations:** Department of Botany and Plant Pathology, Oregon State University, Corvallis, OR, USA; Center for Genome Research and Biocomputing, Oregon State University, Corvallis, OR USA

**Keywords:** *Pseudomonas syringae*, GacA, virulence, two-component response regulator

## Abstract

*Pseudomonas syringae* is a taxon of plant pathogenic bacteria that can colonize and proliferate within the interior space of leaf tissue. This process requires *P. syringae* to rapidly upregulate the production of virulence factors including a type III secretion system (T3SS) that suppress host defenses. GacS/A is a two-component system that regulates virulence of many plant and animal pathogenic bacteria including *P. syringae*. We recently investigated the virulence defect of strain AC811, a Tn*5*::*gacA* mutant of *P. syringa*e pv. *tomato* DC3000 that is less virulent on Arabidopsis. We discovered that decreased virulence of AC811 is not caused by loss of GacA function. Here, we report the molecular basis of the virulence defect of AC811. We show that AC811 possesses a nonsense mutation in *anmK*, a gene predicted to encode a 1,6-anhydromuramic acid kinase involved in cell wall recycling. Expression of a wild-type allele of *anmK* partially increased growth of AC811 in Arabidopsis leaves. In addition to the defective *anmK* allele, we also show that the Tn*5* insertion in *gacA* exerts a polar effect on *uvrC*, a downstream gene encoding a regulator of DNA damage repair. Expression of the wild-type *anmK* allele together with increased expression of *uvrC* fully restored the virulence of AC811 during infection of Arabidopsis. These results demonstrate that defects in *anmK* and *uvrC* are together sufficient to account for the decreased virulence of AC811, and suggest caution is warranted in assigning phenotypes to GacA function based on insertional mutagenesis of the *gacA*-*uvrC* locus.

## INTRODUCTION

*Pseudomonas syringae* is a plant pathogenic bacterium that infects a broad range of host species including numerous agronomically-important food crops and ornamental plants. Due to its genetic tractability, well-defined infection cycle, and ability to infect model plants such as Arabidopsis, *P. syringae* has been adopted as a model organism for studying the genetic bases of bacterial pathogenesis [1]. *P. syringae* can infect most aerial plant tissues, but is commonly studied for its ability to colonize and infect leaves. Upon introduction to a plant host, *P. syringae* is capable of persisting as an epiphyte on leaf surfaces [2]. Under favorable environmental conditions, *P. syringae* swims into the apoplast, or interstitial space, of leaf tissue through stomata or wounds in the leaf surface. Once in the leaf interior, *P. syringae* rapidly proliferates to a high density and causes visible chlorosis as well as water-soaked necrotic lesions on infected tissue.

In order to colonize leaf tissue and cause disease, *P. syringae* must overcome both preformed and induced plant host defenses. A primary line of defense encountered by *P. syringae* are pattern recognition receptors (PRRs) on the surface of plant cells [3]. PRRs monitor the extracellular space for the presence of conserved microbial features and, upon activation, rapidly initiate defense responses to limit pathogen growth. To counteract PRR-activated defenses, *P. syringae* deploys the type III secretion system (T3SS), a syringe-like translocon that delivers bacterial effector proteins directly into plant cells. Within the host cytosol, many of these effector proteins inhibit PRRs or their downstream signaling cascades, thereby suppressing host defenses and promoting *P. syringae* growth within the apoplast [3]. Mutant strains of *P. syringae* that lack a functional T3SS are unable to grow to high levels in host tissues and fail to cause disease.

Although the T3SS is an essential component of *P. syringae* virulence, other factors are also necessary for full infection. These T3SS-independent factors include small molecule toxins that disrupt host immunity [1], enzymes that detoxify plant defense compounds [4], and extracellular proteases [5], and catalases [6] that modify the apoplast environment to enable bacterial survival. Physiological attributes, such as flagella and pili that facilitate bacterial motility, have also been shown to play important roles in the progression of infection [7-8]. Furthermore, signaling pathways that coordinate the expression of these diverse virulence factors are also fundamental to the development of bacterial plant disease [9]. Recent analyses of the *P. syringae* transcriptome during host infection identified various other metabolic, transport, and abiotic stress pathways that are differentially expressed throughout various stages of plant infection [10-12] and that may potentially contribute to pathogenesis. Given the complex, multigene process of adapting to the host environment, it is likely that many additional unknown genetic factors are required for *P. syringae* to successfully grow within the apoplast and cause disease.

GacS/A is a highly conserved two component system that is hypothesized to function as a global regulator of diverse cellular processes including motility, exopolysaccharide biosynthesis, exoenzyme secretion, and T3SS deployment [13-15]. In *Pseudomonas syringae* pv. *tomato* DC3000, a model pathogen capable of infecting both Arabidopsis and tomato, GacS/A has been characterized as a positive regulator of T3SS expression and virulence. Predictive models were developed primarily on the basis of characterizations of a single mutant strain AC811 that contains a Tn*5* transposon insertion within the *gacA* open reading frame [16]. In a recent study, we re-examined phenotypes of AC811 and discovered that, contrary to previous reports, this mutant strain has increased, rather than decreased, expression of T3SS genes during infection of Arabidopsis [17]. Furthermore, we demonstrated that decreased growth of AC811 within Arabidopsis leaves was not due to loss of GacA function. Together, these data indicated that AC811 possesses a T3SS- and GacA-independent defect that compromises its virulence.

In this work we investigated the genetic basis of decreased virulence of strain AC811. We demonstrate that a defective allele of *anmK*, as well as a polar effect of Tn*5* insertion on *uvrC* expression, are sufficient to account for the decreased growth of AC811 in Arabidopsis leaves. Together, these data demonstrate that *anmK* and *uvrC* are required for maximum virulence of DC3000 in Arabidopsis. In addition to virulence phenotypes, we also show that *uvrC*, but not *gacA*, is required for siderophore production by DC3000. These findings highlight potential challenges with assigning functions to GacA based solely on analysis of transposon insertional mutants, and underscore the importance of complementation of GacA-associated phenotypes.

## RESULTS

### Decreased relative growth of AC811 occurs in Arabidopsis leaves but not in King’s B medium

AC811 is unable to grow to high levels or cause disease to Arabidopsis [16-17]. To determine whether AC811 has a general growth defect, we cultured DC3000, AC811 and a DC3000 Δ*gacA* deletion strain in King’s B (KB) broth at 28°C, a temperature frequently used for overnight culturing of *P. syringae*. At 28°C, growth rates of AC811 and DC3000 Δ*gacA* were not significantly different from that of DC3000 (Fig. S1A). We also tested growth of these strains in KB broth at 21°C, the same temperature we use to maintain Arabidopsis plants during infection assays, and observed no growth defect for either AC811 or DC3000 Δ*gacA* (Fig. S1B). These data indicate that AC811 does not have a general growth defect, and suggest that growth of this strain may be specifically hindered within the leaf apoplast environment.

### A nonsense mutation in *anmK* contributes to decreased virulence of AC811

Decreased virulence of AC811 in the apoplast of Arabidopsis leaves is not due to loss of *gacA* [17]. We hypothesized that another mutation may be responsible for the decreased virulence of AC811. We used the Illumina HiSeq platform to sequence the genomes of strain AC811 and its DC3000 parental strain, enabling us to search for single nucleotide polymorphisms (SNPs) that were unique to AC811 [18]. We identified a total of 120 high confidence SNPs in AC811 relative to the published DC3000 reference genome (Table S1). A total of 77 of these SNPs are unique to AC811, as determined based on their absence in the genome sequence of the parental strain. Of these, two SNPs were identified as the most likely to significantly impact protein function and cause the AC811 mutant phenotype. One of these unique SNPs results in a nonsense mutation in the first third of the coding region of PSPTO_0606, encoding a predicted 1,6-anhydromuramic acid kinase (AnmK). We sequenced the *anmK* locus of AC811 and its DC3000 parental strain, confirming that the nonsense mutation is present in AC811 but absent in the parental strain (Fig. 1A). We also identified a SNP exclusive to AC811 that causes a frameshift in PSPTO_1067, a predicted glycosyltransferase. All other SNPs unique to AC811 were either predicted missense mutations, synonymous mutations, or mutations within intergenic regions, and were not predicted to have severe effects on protein function (Table S1).

**Fig 1.**
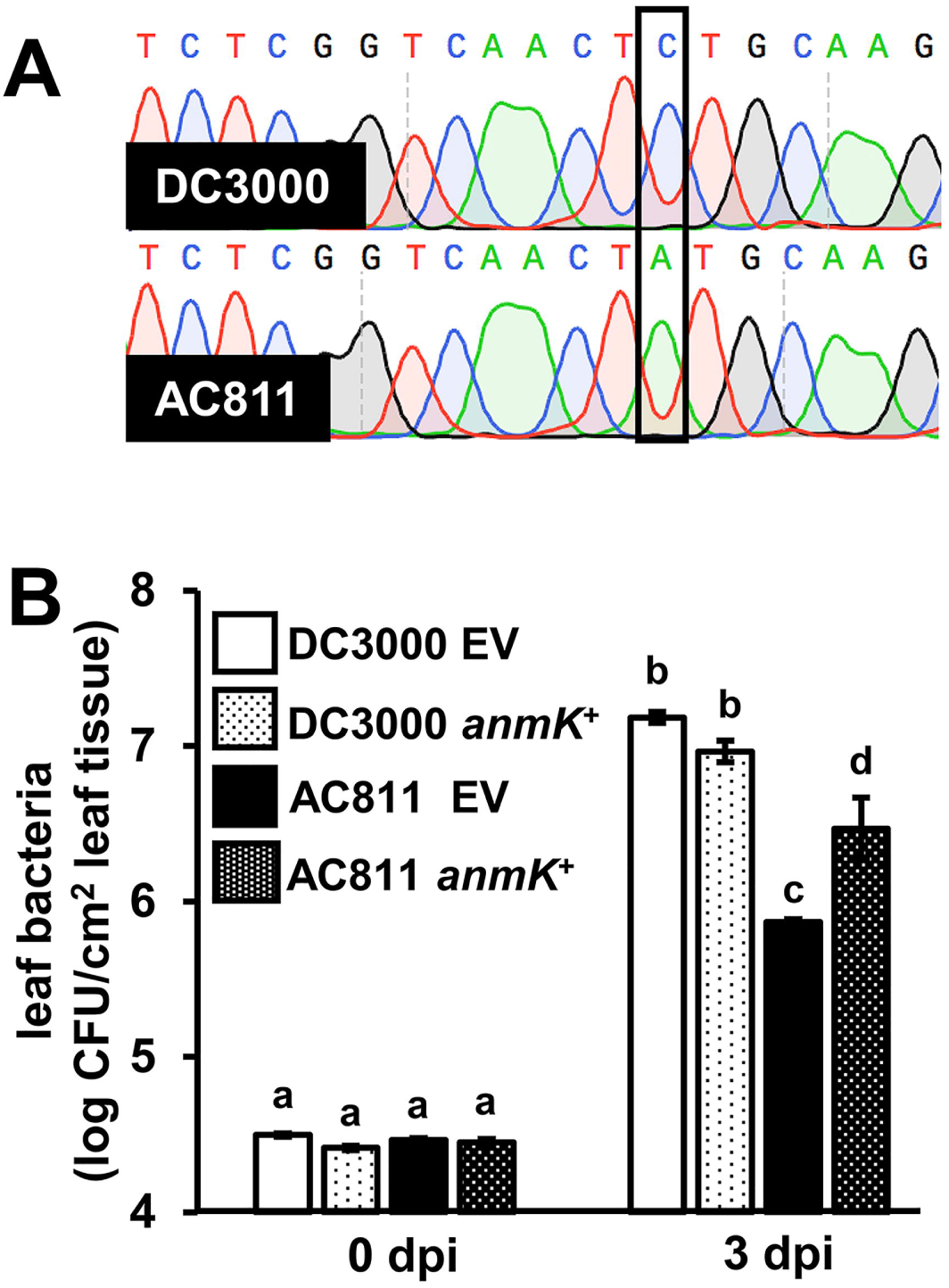
A nonsense mutation in *anmK* contributes to decreased growth of AC811 in Arabidopsis leaves. **(A)** Sanger sequencing chromatograms of the *PSPTO_0606* locus PCR-amplified from genomic DNA of DC3000 and AC811. **(B)** Leaves of Col-0 *Arabidopsis* plants were syringe-infiltrated with DC3000 or AC811 carrying either empty pME6010 (EV) or pME6010::*anmK* plasmid. Data shown are means ± SE of leaf bacteria measured by serial dilution plating. Abbreviation dpi is days post-infection. Statistical groups were determined by ANOVA with multiple pairwise *t-*test comparisons and Tukey’s post-hoc HSD analysis, *p* < 0.05. Data are representative of three independent experiments.

To assess whether the mutations in *anmK* and PSPTO_1067 contribute to loss of AC811 virulence, we individually cloned the DC3000 alleles of *anmK* and PSPTO_1067 into pME6010, introduced the resulting constructs into DC3000 and AC811, and infected Arabidopsis leaves with these strains by syringe-infiltration. AC811 carrying *anmK*::pME6010 grew to significantly higher levels relative to AC811, indicating that introduction of the DC3000 allele of *anmK* partially complemented the ability of AC811 to grow *in planta* (Fig. 1B). However, growth of AC811 *anmK*::pME6010 was not fully restored to DC3000 levels, suggesting that other factors may contribute to AC811 *in planta* growth (Fig. 1B). Expression of *anmK* did not significantly impact the growth of wild type DC3000 (Fig. 1B). DC3000 is a highly virulent pathogen that can achieve high levels of growth, which could mask any potential growth-promoting effects of *anmK*. To test this, we introduced *anmK*::pME6010 into a virulence-attenuated DC3000 Δ*avrPto*Δ*avrPtoB* strain [19]. Similar to DC3000, we detected no *anmK*-dependent increase in the growth of this strain in Arabidopsis leaves (Fig. S2). No increase in growth was measured for AC811 carrying *PSPTO_1067*::pME6010, suggesting that the frameshift mutation of this gene does not negatively impact AC811 virulence (Fig. S3). Together, these data indicate that a nonsense mutation in *anmK* contributes to the decreased virulence of AC811.

### Tn*5* insertion in *gacA* disrupts the expression of downstream *uvrC*

We reasoned that the insertion of Tn*5* into the *gacA* open reading frame in AC811 may have a polar effect on nearby genes. In DC3000, *gacA* is predicted to be part of an operon including *uvrC* and *pgsA* [13] that is conserved among γ-proteobacteria [14-15, 20-22] (Fig. 2A). The *uvrC* gene encodes a component of the dinucleotide excision repair (NER) complex involved in remediating DNA damage [23], whereas *pgsA* is predicted to encode a protein involved in phospholipid biosynthesis [13]. In *E. coli*, a similarly structured operon can be transcribed either as a polycistronic transcript, or as separate monocistronic transcripts due to promoter elements located within the *gacA* open reading frame [24].

**Fig 2.**
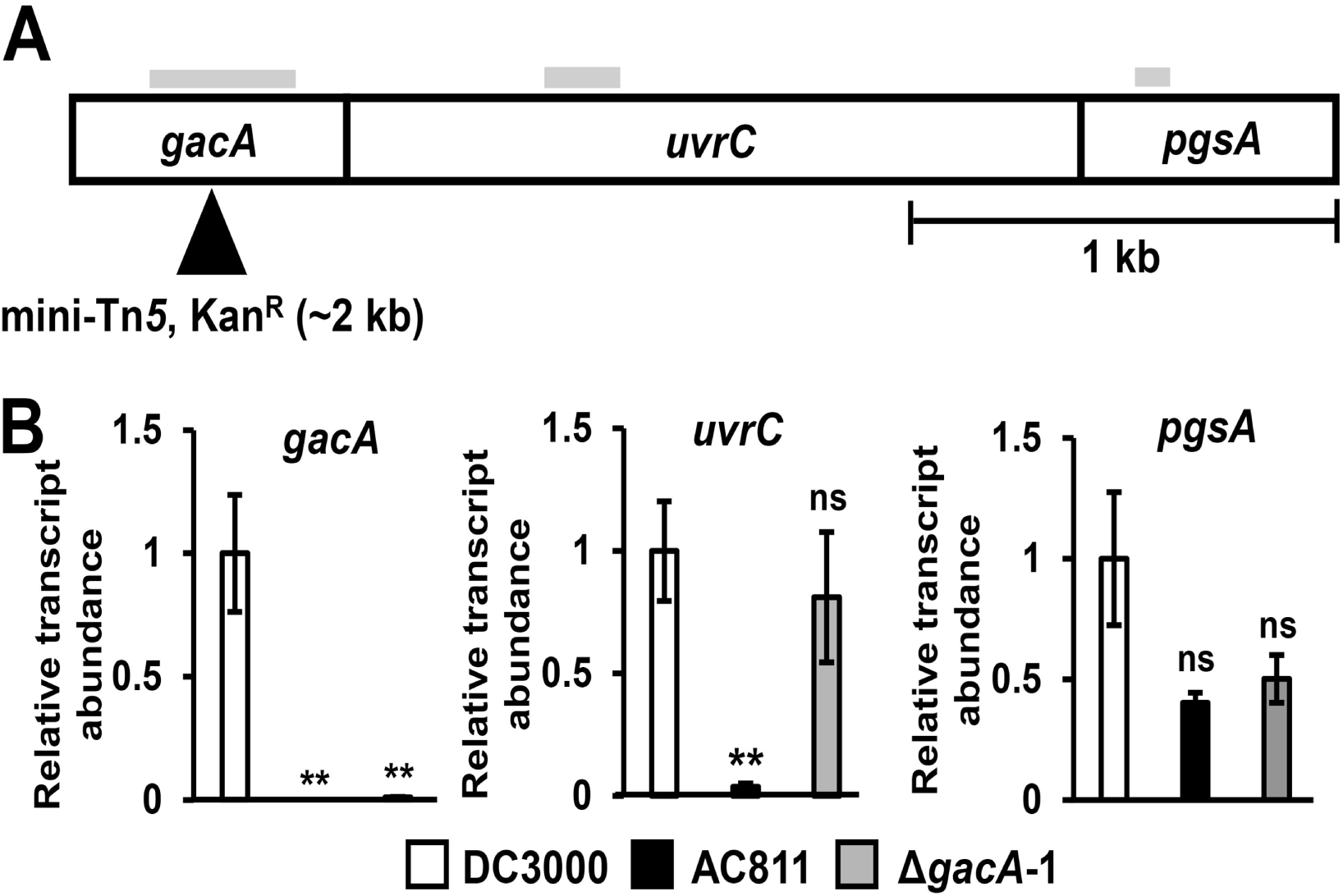
Expression of *uvrC* is decreased in AC811 but not in Δ*gacA*-1. **(A)** Schematic of the putative *gacA-uvrC-pgsA* operon in DC3000. Filled triangle shows mini-Tn*5* insertion site in AC811. Solid black vertical bars indicate predicted translation start sites of *uvrC* and *pgsA*. Shading above each gene indicates regions of mRNA transcripts amplified by qRT-PCR. **(B)** Quantitative RT-PCR analysis of *gacA, uvrC*, and *pgsA* transcripts. Transcript abundances were normalized to *gyrA* transcripts using the PCR efficiency ^−ΔΔC□t^ method and calculated relative to transcript levels measured in wild type DC3000. Graphed are means ± SE from data pooled across two independent experiments with two technical replicates each; n = 4. Asterisks denote statistical significance from pairwise *t*-tests performed between the indicated strains and DC3000. \*\**p* < 0.01; ns = no significant difference (*p* > 0.05).

We used quantitative RT-PCR (qRT-PCR) and gene-specific primers to measure the abundance of *gacA*-, *uvrC*- and *pgsA*-containing transcripts in AC811. As expected, we could not detect *gacA*-containing transcripts from AC811 or the DC3000 Δ*gacA*-1 deletion strain (Fig. 2B). Transcripts containing *uvrC* were also significantly decreased in AC811 bacteria relative to levels in DC3000 (Fig. 2B). In contrast, the abundance of *pgsA*-containing transcripts was not significantly altered in AC811 (Fig. 2B). We also measured *uvrC* and *pgsA* expression in DC3000 Δ*gacA*-1 and found that the abundance of *uvrC* transcripts was similar to DC3000 levels, whereas the average abundance of *psgA* transcripts was reduced approximately 2-fold, although this reduction was not significantly different from *psgA* levels in either DC3000 or AC811 (Fig. 2B). For all genes assessed, virtually no transcripts were detected from negative control reactions in which cDNA was mock synthesized in the absence of reverse transcriptase (Fig. S4). Based on these data, we conclude that the Tn*5* insertion in *gacA* decreases the expression of *uvrC* but not *pgsA*. Furthermore, these data indicate that deletion of the entire *gacA* open reading frame does not exert a significant polar effect on the expression of downstream genes.

We additionally assessed whether polycistronic transcripts containing both *gacA* and *uvrC* are altered in AC811. To investigate, we performed qRT-PCR with primers that span the *gacA*-*uvrC* junction and measured a significant decrease in these transcripts in AC811 (Fig. S5). We also detected transcripts containing the *uvrC*-*pgsA* junction by qRT-PCR, but did not observe any significant change in the abundance of these transcripts in either AC811 or DC3000 Δ*gacA*-1 (Fig. S5B).

### Expression of *uvrC* and *anmK* fully rescues the growth defect of AC811 in Arabidopsis

To determine whether decreased *uvrC* expression contributes to the virulence defect of AC811, we cloned *uvrC* into broad host range vector pME6010 and introduced this construct into both DC3000 and AC811. We then syringe-infiltrated theses strains into Arabidopsis leaves and observed that AC811 carrying *uvrC*::pME6010 grew to significantly higher levels than AC811 carrying empty pME6010 (Fig. 3A). Similar to our results with *anmK* expression, *uvrC*::pME6010 did not alter growth of DC3000 WT (Fig. 3A) or DC3000 Δ*avrPto*Δ*avrPtoB* (Fig. S2) in Arabidopsis. To alleviate the polar effect of Tn*5* on *uvrC*, we used allelic exchange to delete both *gacA* and Tn*5* from the AC811 genome, generating an AC811 Δ*gacA* strain. Expression of *uvrC* in AC811 Δ*gacA* was restored to DC3000 levels as determined by qRT-PCR (Fig. S5C). Growth of AC811 Δ*gacA* was significantly higher relative to AC811 (Fig. 3B) but was not fully restored to DC3000 or DC3000 Δ*gacA* levels.

**Fig 3.**
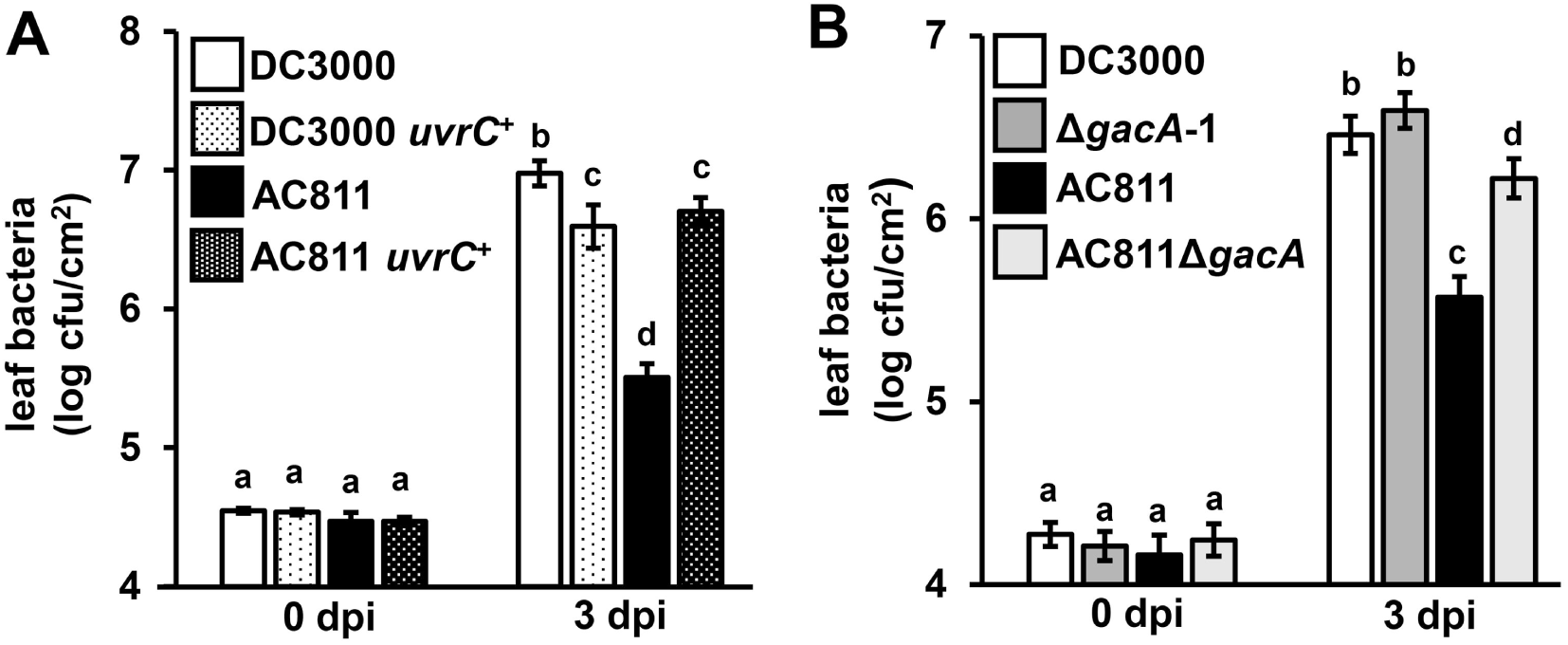
Growth of AC811 in Arabidopsis is enhanced by eliminating polar effects of Tn*5*:*gacA* insertion on *uvrC* expression. **(A)** Arabidopsis leaves were syringe-infiltrated with DC3000 or AC811 carrying either pME6010 empty vector or pME6010::*uvrC* as indicated. Graphed are means ± SE of leaf bacteria measured by serial dilution plating; n = 9. Data are pooled from three independent experiments. **(B)** Arabidopsis leaves were syringe infiltrated with DC3000 WT, AC811, Δ*gacA*-1 or AC811 Δ*gacA*. Graphed are means ± SE of leaf bacteria measured by serial dilution plating; n = 14. Abbreviation dpi is days post-infection. Data are pooled from four independent experiments. Small-case letters in (A) and (B) denote statistical groups determined by ANOVA with multiple pairwise *t*-test comparisons and Tukey’s post-hoc HSD analysis, *p* < 0.05.

We next introduced a plasmid with the DC3000 allele of *anmK* into AC811 Δ*gacA*. The introduction of *anmK*_DC3000_ was sufficient to fully restore the growth of AC811 Δ*gacA* to DC3000 levels in Arabidopsis leaves (Fig. 4). Disease symptoms caused by AC811 Δ*gacA* carrying *anmK*_DC3000_, such as chlorosis of infected leaves, also appeared similar to those produced by DC3000 infection (Fig. S6). Based on these data we conclude that decreased *uvrC* expression and loss of AnmK function are together causal for decreased virulence of AC811 in Arabidopsis leaves.

**Fig 4.**
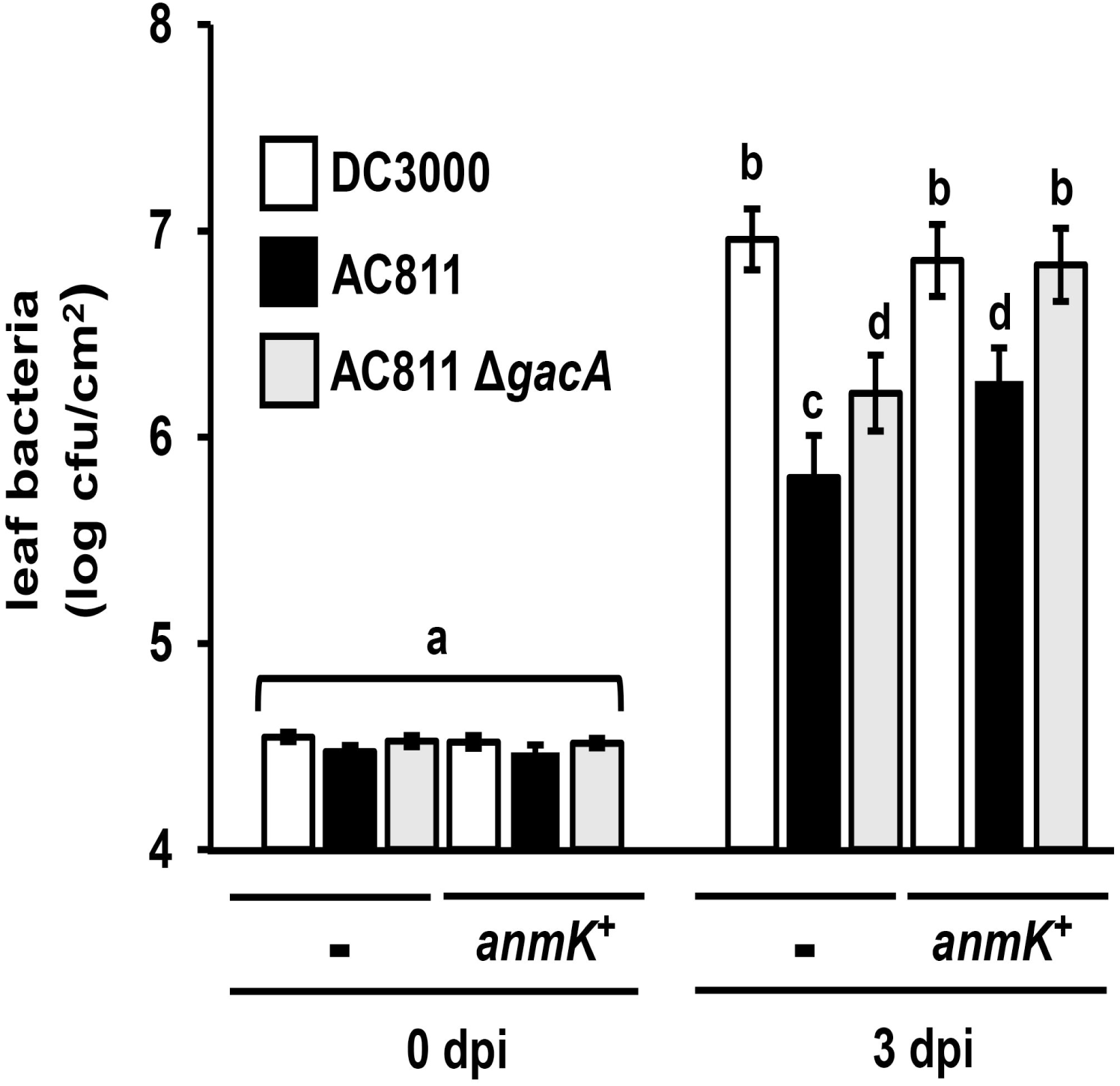
Growth of AC811 in Arabidopsis is fully restored by eliminating Tn*5* effects on *uvrC* and by expression of a wild-type *anmK* allele. Arabidopsis leaves were syringe-infiltrated with DC3000, AC811 or AC811 Δ*gacA* carrying either pME6010 empty vector or pME6010::*anmK*. Graphed are means ± SE of leaf bacteria enumerated by serial dilution plating; n = 3. Abbreviation dpi is days post-infection. Data are pooled from four independent experiments; n = 12. Statistical groups were determined by ANOVA with multiple pairwise *t*-test comparisons and Tukey’s post-hoc HSD analysis, *p* < 0.05.

### Decreased *uvrC* expression reduces siderophore production in DC3000

The biosynthesis and extracellular secretion of iron-chelating siderophores is positively regulated by GacS/A in certain pseudomonads [25], but suppressed by GacS/A in others [26-27]. To investigate if GacA regulates siderophore production in DC3000, we incubated DC3000, AC811 and DC3000 Δ*gacA* on CAS blue agar to detect siderophore production. We observed a significant reduction in siderophore production by AC811 (Fig. 5A,B). However, no decrease in siderophore levels was observed for DC3000 Δ*gacA*, indicating the decrease in siderophore production by AC811 is *gacA*-independent (Fig. 5A,B). Expression of *uvrC* in AC811 significantly increased siderophore production by AC811 to near DC3000 levels (Fig. 5C). Furthermore, siderophore production by AC811 Δ*gacA* was indistinguishable from DC3000 (Fig. 5B). Together, these data indicate that, similar to DC3000 growth in the leaf apoplast, polar effects of Tn*5* on *uvrC* expression influence the levels of siderophore production.

**Fig 5.**
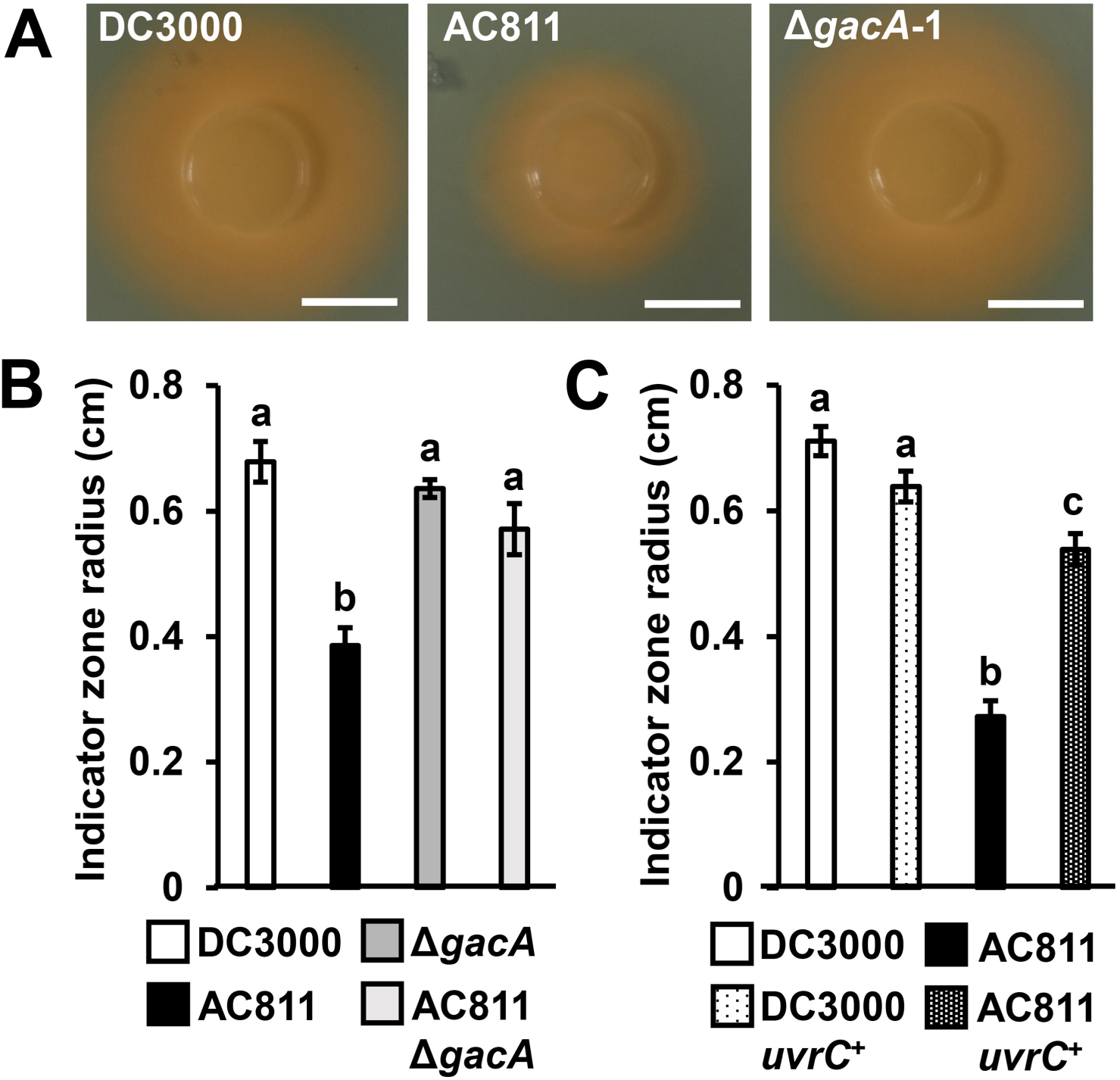
Siderophore production by DC3000 is *gacA*-independent. Siderophore production of DC3000, AC811, and Δ*gacA* was detected on CAS blue agar plates. **(A)** Photographs of agar plates showing fluorescent siderophore production, visible as an orange halo surrounding the bacterial colony. White bar is scale of 1 cm. Results are representative of three independent experiments. **(B)** Radii of indicator zone from strains carrying either pME6010 empty vector or pME6010::*uvrC*. Graphed are means ± SE of data pooled from two independent experiments; n = 7. **(C)** Radii (cm) of the zone of indicator color change were measured from the edge of bacterial colony. Graphed are means ± SE of data pooled from three independent experiments; n = 9. Small case letters in (B) and (C) denote statistical groupings determined by ANOVA with multiple pairwise *t*-test comparisons and Tukey’s post-hoc HSD analysis, *p* < 0.05.

## DISCUSSION

Isolating and characterizing mutants that have loss-of-virulence phenotypes is fundamental to understanding virulence mechanisms of *P. syringae* and other bacterial pathogens. However, laboratory manipulation of bacterial genomes can present various challenges. Common techniques used to generate mutants, such as Tn*5* insertional mutagenesis, carry the risk of disrupting expression of surrounding genes. Furthermore, accumulation of SNPs or other mutations within the genomes of laboratory strains may further confound the analysis of mutant phenotypes. Here, we investigated the genetic defects of AC811 and demonstrated that a polar effect of the Tn*5* transposon, as well as a second-site mutation are causal for decreased virulence of AC811. In this case, these unintended genetic defects led to the discovery of two genes that positively regulate DC3000 virulence. However, these *gacA*-independent effects also contributed to confusion regarding the role of *gacA* in DC3000 virulence, and highlight the need for caution when interpreting phenotypes of mutant strains, as well as the need to confirm that the casual lesion is indeed within the gene of interest. In this work, whole genome sequencing of AC811 proved to be an effective approach for identifying an off-target genetic defect. With the ever-decreasing cost of high-throughput DNA sequencing, we anticipate that whole genome sequencing of mutant strains may become a routine and necessary practice in the laboratory for confirming the results of bacterial genetic studies.

We demonstrated that a lesion in *anmK* (PSPTO_0606) causes a reduction in *in planta* growth of AC811. The *anmK* gene is predicted to encode for anhydro-1,6-muramic acid kinase, an enzyme that catalyzes the phosphorylation of muropeptides generated during the natural recycling of bacterial cell wall peptidoglycan [28]. AnmK has not been previously identified as necessary for virulence of *P. syringae*, and the specific role of this enzyme during host infection remains unknown. In *E. coli*, loss of AnmK function is known to disable phosphorylation-dependent retention of muropeptides in the bacterial cytosol, preventing the metabolic reuptake of peptidoglycan. This disruption of cell wall regeneration leads to the extracellular accumulation of murein breakdown products [28]. Among Gram negative bacteria, fluctuations in extracellular muropeptide composition act as signals of environmental stress, such as high cellular osmolarity induced by water stress or cell envelope damage inflicted by antibiotics [29]. AnmK-mediated cell wall recycling could be required for the rapid proliferation of DC3000 following colonization of the host apoplast. Alternatively, abnormal accumulation of muropeptides in the extracellular space may disrupt DC3000 cellular stress signaling, possibly affecting the ability of DC3000 to tolerate stressors of the plant environment such as immune defenses and/or water stress.

Our results suggest that UvrC may play a role in regulating DC3000 virulence. As a subunit of the conserved dinucleotide excision repair (NER) complex, UvrC catalyzes the excision of DNA lesions, contributing to bacterial survival under UV- or mutagen-induced stress [30]. Accordingly, *gacA* mutants in various bacterial species show UV-hypersensitivity, likely due to negative polar effects on *uvrC* expression [15, 20, 22, 31-33]. While UV stress tolerance is known to impact epiphytic survival in *Pseudomonas* spp. [34]. it is unknown how UvrC may contribute to growth of bacteria such as DC3000 that are primarily endophytic and are unable to persist epiphytically on the leaf surface. Similar to our findings, a study of both polar and nonpolar *gacA*^−^ mutants in the animal pathogen *Pseudomonas aeruginosa* revealed that negative effects on *uvrC* expression, rather than inactivation of *gacA*, were associated with a loss of virulence in *Bombyx mori* larvae [33, 35]. Furthermore, an *uvrC* mutant of the human pathogen *Yersinia enterocolitica* has been shown to cause reduced expression of *inv*, a gene required for colonization of the intestinal lymphoid tissues, although possible polar effects on *gacA* expression in this mutant could not be ruled out [36]. Interestingly, normal *inv* expression by the *Y. enterocolitica uvrC* mutant could be restored by expression of either SspA or ClpB, both of which function as regulators of stress tolerance in *E. coli* [32-35]. These results suggest that UvrC may broadly contribute to virulence in a range of pathogens, potentially through an unknown role in the cellular stress response. In DC3000, UvrC may regulate tolerance of environmental stressors encountered in the leaf apoplast enabling bacterial survival and proliferation in host tissues.

Throughout the γ-proteobacteria, *gacA* is located in an operon with *uvrC* and *pgsA*. In numerous species, the translation start site of *uvrC* overlaps the stop codon of *gacA* [21-22, 31, 37]. Previously, *gacA* was reported to be expressed as a single gene in DC3000 based on detection of a ∼0.75 kb transcript by northern blot with a *gacA* sequence-specific probe [16]. Our qRT-PCR data demonstrate that transcripts containing *gacA-uvrC* and *uvrC*-*psgA* junctions are also produced in DC3000, indicating that *gacA* is expressed as part of a larger operon structure than previously reported. Consistent with this operon model of *gacA* and *uvrC* expression, we observed a significant reduction in *uvrC* and *gacA*-*uvrC* transcripts in AC811 due to polar effects of the Tn*5* insertion in *gacA*. Although we also detected bicistronic *uvrC*-*psgA* transcripts, both *psgA*- and *uvrC*-*psgA*-containing transcripts were not significantly altered in abundance in AC811 (Fig. 4 and Fig. S4). In *E. coli*, monocistronic *uvrC* transcripts and bicistronic *uvrC*-*psgA* transcripts are produced by promoter sequences within the *gacA* coding region [24] and these promoter elements are conserved within DC3000 *gacA* based on our sequence analysis. Therefore, *uvrC*-*psgA* transcription in AC811 may occur through promoter elements in *gacA* that are downstream of Tn*5* insertion, allowing for expression of *psgA* to remain unperturbed. Notably, expression of *uvrC* was restored to wild type levels in our AC811 Δ*gacA* strain, indicating that promoter elements upstream of *gacA* are sufficient to drive the expression of *uvrC* even in the absence of *gacA*. Despite the conserved genomic arrangement of *gacA* and *uvrC*, the functional relationship between these genes (if any) remains unknown. Further analysis of the mechanistic role of UvrC during *P. syringae* infection may provide important insights into the functional significance of the *gacA-uvrC* association.

Our results also uncovered a role for *uvrC* in siderophore production by DC3000. Siderophores are a class of fluorescent iron-chelating compounds that are secreted by bacteria into the extracellular space during iron-limiting growth conditions. Upregulation of *uvrC* expression under iron-limiting conditions has been reported in certain bacterial species [38], although this effect is not observed in all cases [39]. Both *uvrC* expression and siderophore production are induced in biofilm-forming cells of *Mycobacterium smegmatis*, though it is unknown whether *uvrC* influences siderophore production or is simply concomitantly expressed [40]. Siderophores contribute to the virulence of many bacterial pathogens. However, a DC3000 mutant that is unable to synthesize or import siderophores remains fully virulent on host tomato plants [41]. Therefore, although both decreased virulence and siderophore production are observed with AC811, it is unlikely that these two phenotypes are causally-related. In addition to identifying UvrC as a regulator of siderophore production, our results also demonstrate that GacA is dispensable for this phenotype. Regulation of siderophore biosynthesis by GacS/A has been described in various Pseudomonads, though whether GacS/A functions as a positive or negative regulator of siderophore production appears to vary across species [25-27]. As many previous studies have relied on insertional mutants to assess the function of GacA, in some cases it may be necessary to re-assess these mutant strains to determine if off-target effects on *uvrC* expression may be responsible for *gacA*-associated phenotypes.

## Materials and Methods

### Bacterial strains and growth conditions

*P. syringae* strains stored at −80°C were streaked from 25% glycerol stocks onto King’s B medium (KBM) 1.5% agar plates. The KBM agar was supplemented with 50 μg/mL rifampicin and antibiotics (10 μg/mL tetracycline, 20 μg/mL gentamycin, 50 μg/mL chloramphenicol) as necessary. For all experiments *P. syringae* were grown at room temperature on KBM agar for 2 days prior to use. *E. coli* strains were grown at 37 °C in Luria Broth (LB) supplemented with antibiotics (5 μg/mL tetracycline, 25 μg/mL gentamycin, 30 μg/mL chloramphenicol) as necessary.

### Construction of *anmK* and *uvrC* complementation plasmids

The Gibson assembly method was used to clone the open reading frames of *uvrC* and *anmK* into broad host range vector pME6010 under control of a constitutive kanamycin promoter [42]. Primers 6010-F1/6010-R1 and 6010-F2/6010-R2 (Table S2) were used to PCR amplify pME6010 as 3.0-kb and 5.5-kb DNA fragments. Primer sets 6010-uvrC-F/6010-uvrC-R and 6010-anmK-F/6010-anmK-R (Table S2) were used to amplify *uvrC* and *anmK*, respectively, from DC3000 genomic DNA. PCR products of *uvrC* or *anmK* were then mixed with the 3.0- and 5.5-kB fragments of pME6010 and assembled using the NEBuilder HiFi mix (NEB). Plasmids were transformed into *E. coli* DH5a competent cells (NEB) by heat shock. Tc^R^ clones were screened for the presence of the *uvrC* or *anmK* insert by colony PCR with primer sets 6010-uvrC-F/6010-uvrC-R or 6010-anmK-F/6010-anmK-R, respectively. Constructs were confirmed by Sanger sequencing. Plasmids were conjugated into DC3000 by tri-parental mating using an *E. coli* helper strain carrying pRK600 [43].

### Generation of *gacA* deletions in DC3000

The previously described Δ*gacA*::pK18*mobsacB* suicide vector [17] was conjugated into DC3000 by tri-parental mating using an *E. coli* helper strain carrying pRK600 [43]. Merodiploid colonies were passaged 1-3 days in KB broth with 50 μg/mL rifampicin prior to counter-selection on KB agar with 15% sucrose. Sucrose-resistant colonies were plated on KB agar with 50 μg/mL rifampicin supplemented with or without selective antibiotics. PCR reactions with primers gacA-F and gacA-R (Table S2) were used to screen genomic DNA isolated from kanamycin-sensitive colonies for deletion of *gacA*. To generate a Δ*gacA* deletion in AC811 (Kan^R^), the Δ*gacA* deletion cassette from pK18*mobsacB* was sub-cloned into pT18*mobsacB* (Addgene #72648, Tc^R^) using the Gibson assembly method. Primers pT18gacA-F and pT18gacA-R (Table S2) were used to PCR amplify the Δ*gacA* deletion cassette from pK18*mobsacB*. Primers pT18-F and pT18-R (Table S2) were used to PCR amplify the entire backbone of pT18*mobsacB*. PCR products were assembled into an intact plasmid using NEBuilder HiFi mix (NEB). Plasmids were transformed into *E. coli* DH5a competent cells (NEB) by heat shock. Clones were screened for the presence of the Δ*gacA* deletion cassette by colony PCR with primers M13F and M13R, and constructs were confirmed by Sanger sequencing.

### Measurement of bacterial growth in culture

*P. syringae* were scraped from agar plates and inoculated into 3 mL cultures of KB broth containing 50 μg/mL rifampicin. Cultures were grown at 28°C overnight in a shaking incubator until cultures reached an optical density at 600 nm (OD_600_) of ∼1.0. Overnight cultures were then diluted to OD_600_ = 0.05 in 30 mL of KB broth containing 50 μg/mL rifampicin in 250-mL volume flasks. Cultures were maintained at 28°C or 21°C with shaking. One mL samples were removed from each flask every 2 hrs for 12 hours and a spectrometer was used to measure the OD600.

### *In planta* bacterial growth assays

*Arabidopsis thaliana* Col-0 seeds were sterilized, stratified and sown onto MS agar plates as described previously [9]. After two weeks of growth, seedling were transplanted to Sunshine mix soil (Sun Gro Horticulture) in 24-well flats. Plants were maintained in a 22°C, 10-hour day growth chamber before and during the infection. To prepare the inoculum, *P. syringae* were scraped from the surface of KB agar and vortexed in 1 mL of sterile H_2_O. Cells were then washed twice and pelleted between each wash by centrifugation at 21,000 × *g* for 1 min, followed by resuspension of bacterial pellets in 1 mL H_2_O. Bacterial suspensions were diluted with H_2_O to an OD_600_ of 0.001 (∼1 × 10^6^ colony-forming units/mL). A needle-less syringe was used to infiltrate bacteria into leaves of five week-old plants. Three to five leaves were infected per plant, and individual plants were infected with only a single strain of bacteria. At times indicated, a cork borer was used to remove 0.2 cm^2^ discs from infected leaves. For each sample, three leaf disks from a single plant were pooled into a single microcentrifuge tube and homogenized in 500 μL H_2_O by grinding the leaf tissue with a plastic pestle. The resulting homogenate was serially-diluted and pipetted in 10 μL volumes onto the surface of KB agar containing 50 μg/mL rifampicin. After 24 hours bacteria, agar plates were viewed under a stereomicroscope to visualize and enumerate individual colonies.

### Genome sequencing and SNP calling

The Qiagen DNeasy Blood & Tissue Kit Genomic DNA was used to extract genomic DNA from DC3000 and AC811. The Nextera XT library prep kit was used to prepare sequencing libraries from each genomic DNA sample. The resulting libraries were sequenced (150 bp paired end) on a single lane of an Illumina HiSeq 3000 instrument at the Center for Genome Research and Biocomputing at Oregon State University. Reads were aligned to the *Pseudomonas syringae* pv. tomato DC3000 reference genome (NCBI: 223283), using bowtie2 version 2.3.2 with the option “—local” [44]. Alignments were converted to bam format using samtools version 0.1.19 and read groups were added using Picard tools version 2.0.1 [45] (Picard Tools, 2015). GATK version 3.7 HaplotypeCaller with the options ‘-ERC GVCF -ploidy 1’ was used to call variants, and the data were then combined using GenotypeGVCFs [46]. The program snpEff version 4.3T was used to predict effects of SNPs on protein function [47].

### Quantification of bacterial gene expression by qRT-PCR

Bacteria were grown overnight at 20°C in KB broth containing 50 μg/mL rifampicin to an OD_600_ = ∼1.0. Cells were centrifuged for 10 min at 21,000 × *g* and the resulting bacterial pellets were frozen in liquid nitrogen and stored at −80°C prior to use. RNA was extracted from the bacterial pellets and 500 ng converted into cDNA in 25 μL reactions as described previously [9]. Quantitative RT-PCR (qRT-PCR) reactions were performed in 10 μL volumes using 5 μL of SsoAdvanced Universal SYBR Green Supermix (Bio-Rad), 4 μL of a mix containing 0.5 μM of each transcript-specific primer (Table S2), and 1 μL of cDNA. SYBR Green fluorescence from each qRT-PCR reaction was monitored using the C1000 Thermal Cycler with CFX96 Real-Time System (Bio-Rad). Relative abundance of transcripts was calculated relative to *gyrA* using the formula Transcript Abundance = PCR efficiency^−(Ct[*gene*]-Ct[*gyrA*])^ [48]. PCR efficiency values were calculated for each qRT-PCR reaction using LinRegPCR [49].

### Detection of siderophore production

Chrome azurol (CAS) blue agar plates [50] containing 2% (w/v) sucrose, 0.2% asparagine, 0.1% KH_2_PO_4_, 0.05% MgSO_4_-7H_2_O, 0.1 M PIPES salt, and 1.5% agar were prepared. *P. syringae* were scraped from the surface of KB agar and suspended in 1 mL of sterile H_2_O by vortexing. Cells were then washed three times with 1 mL of sterile H_2_O. Bacterial suspensions were diluted with H_2_O to an OD_600_ of 1.0 and 5 μL spotted at the center of each plate. Plates were incubated in the dark at room temperature for 48 hr prior to imaging. Radii of the diffuse orange halos that appear in the agar were measured starting from the edge of the bacterial colony.

## ACKNOWLEDGMENTS

We thank the Department of Botany and Plant Pathology at Oregon State University for its generous support of the computing cluster.

## SUPPORTING INFORMATION

**Table S1. Single nucleotide polymorphisms identified by Illumina sequencing.**

**Table S2. Sequences of oligonucleotide primers used in this study.**

**Table S3. List of strains used in this study.**

**Fig S1.**
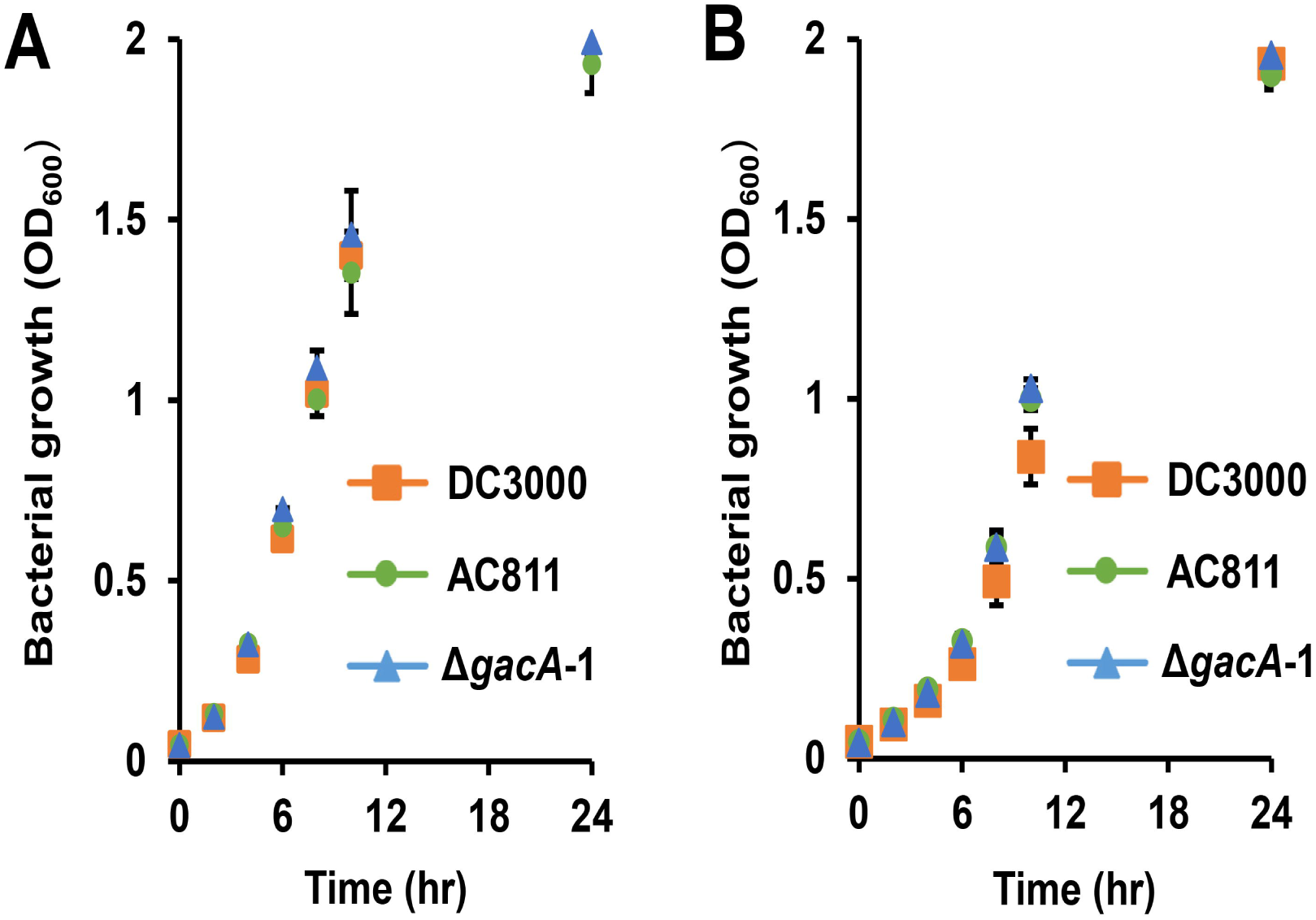
*gacA* mutants exhibit wild type levels of growth in culture. Time-course of bacterial growth at 28°C. DC3000, AC811 and Δ*gacA*-1 were inoculated at OD_600_ = 0.05 into KB broth and incubated at **(A)** 28°C or **(B)** 21°C with shaking. Graphed are means ± SE of OD_600_, n = 3. Data are pooled from three independent experiments.

**Fig S2.**
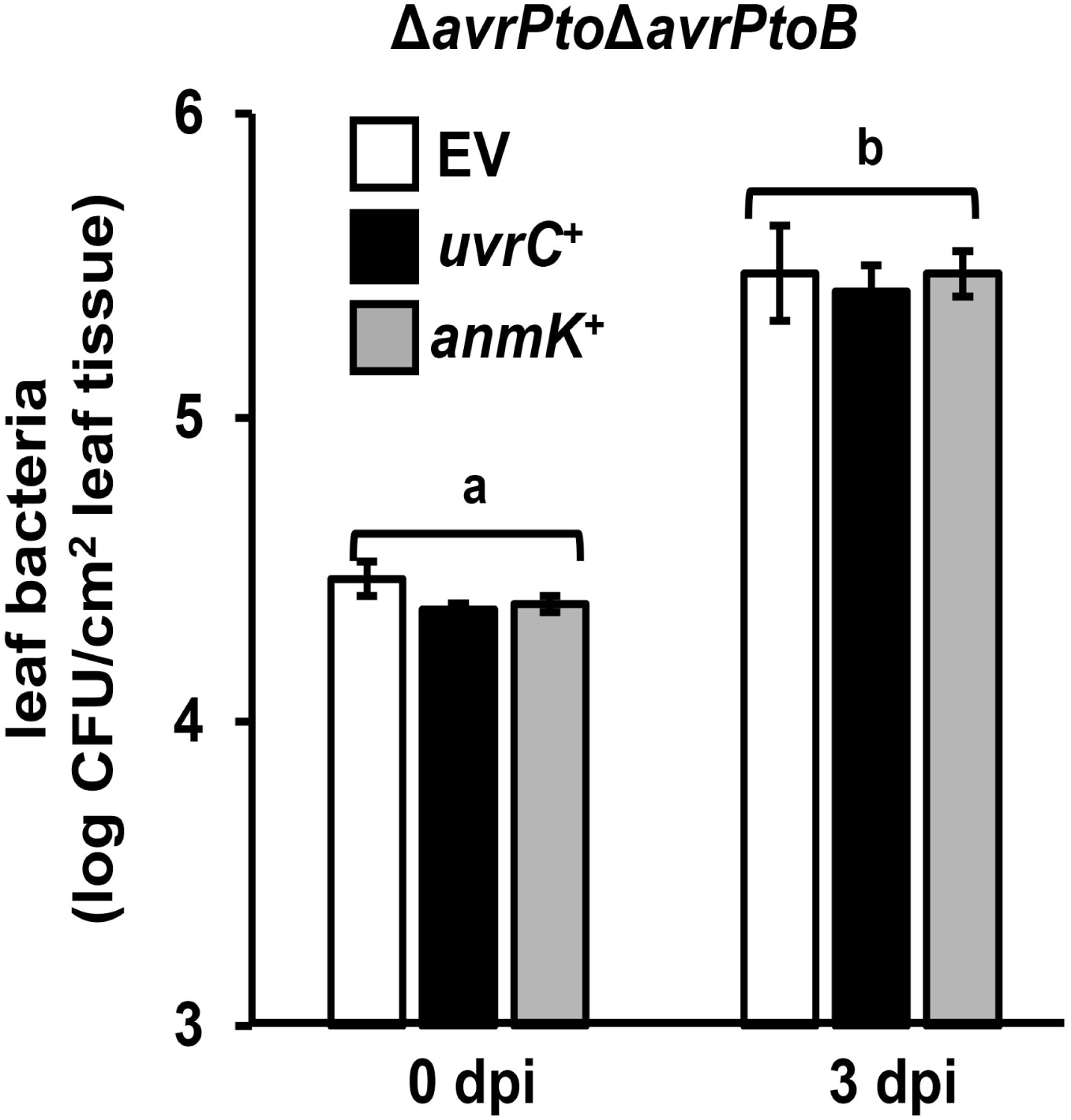
Expression of *uvrC* or *anmK* does not alter growth of DC3000 Δ*avrPto*Δ*avrPtoB.* Arabidopsis leaves were syringe-infiltrated with DC3000 Δ*avrPto*Δ*avrPtoB* carrying empty pME6010 (EV), pME6010::*uvrC*, or pME6010::*anmK* as indicated. Abbreviation dpi is days post-infection. Graphed are means ± SE from data pooled from two independent experiments; n = 6.

**Fig S3.**
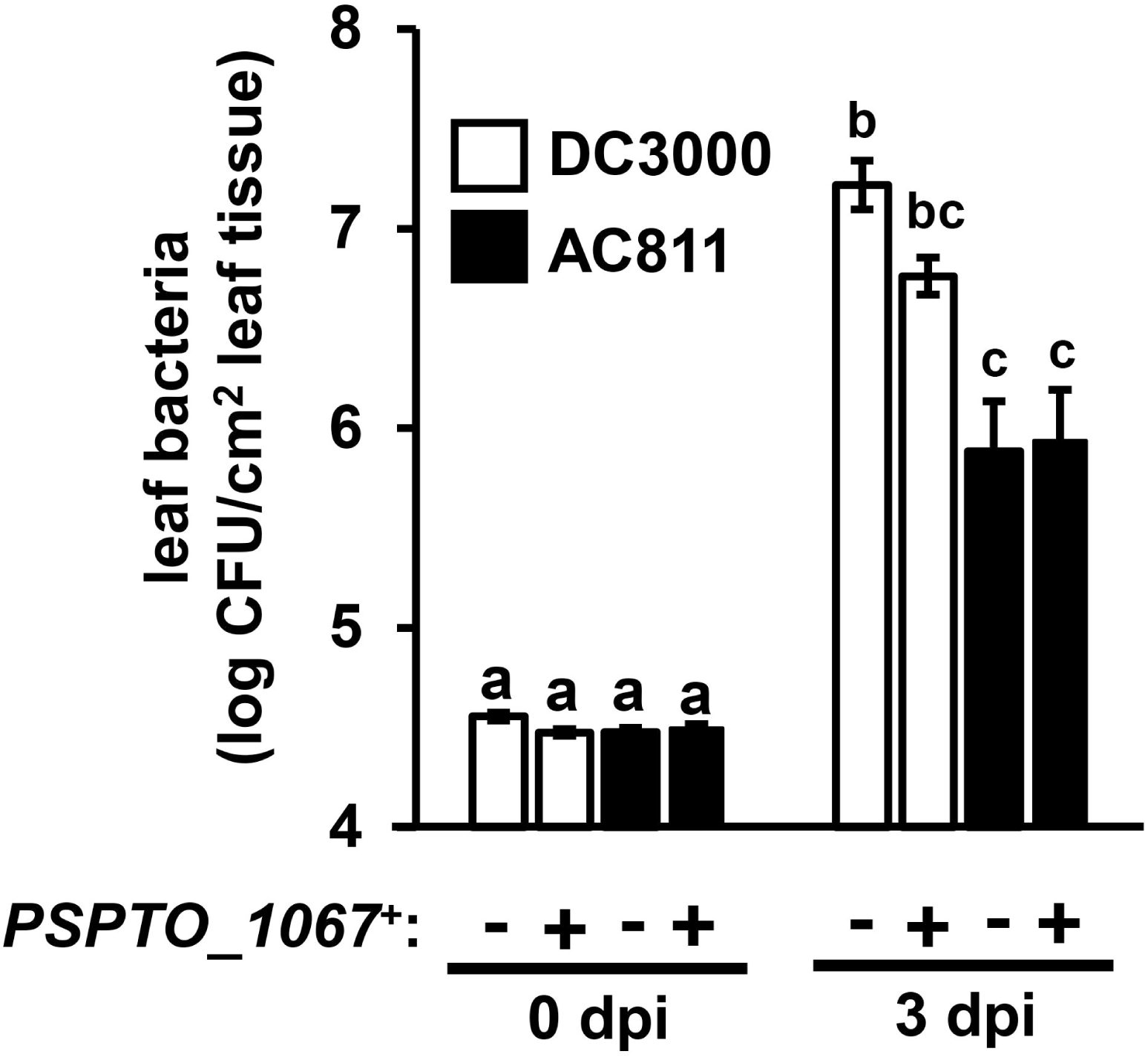
Expression of *PSPTO_1067* does not increase growth of AC811. Arabidopsis leaves were syringe infiltrated with DC3000 and AC811 carrying either pME6010 empty vector or pME6010::*PSPTO_1067* plasmid. Graphed are means ± SE of bacterial growth as determined by serial dilution plating, n = 3. Small-case letters denote statistical groups determined by ANOVA with multiple pairwise *t*-test comparisons and Tukey’s post-hoc HSD analysis, *p* < 0.05. Data are representative of two independent experiments.

**Fig S4.**
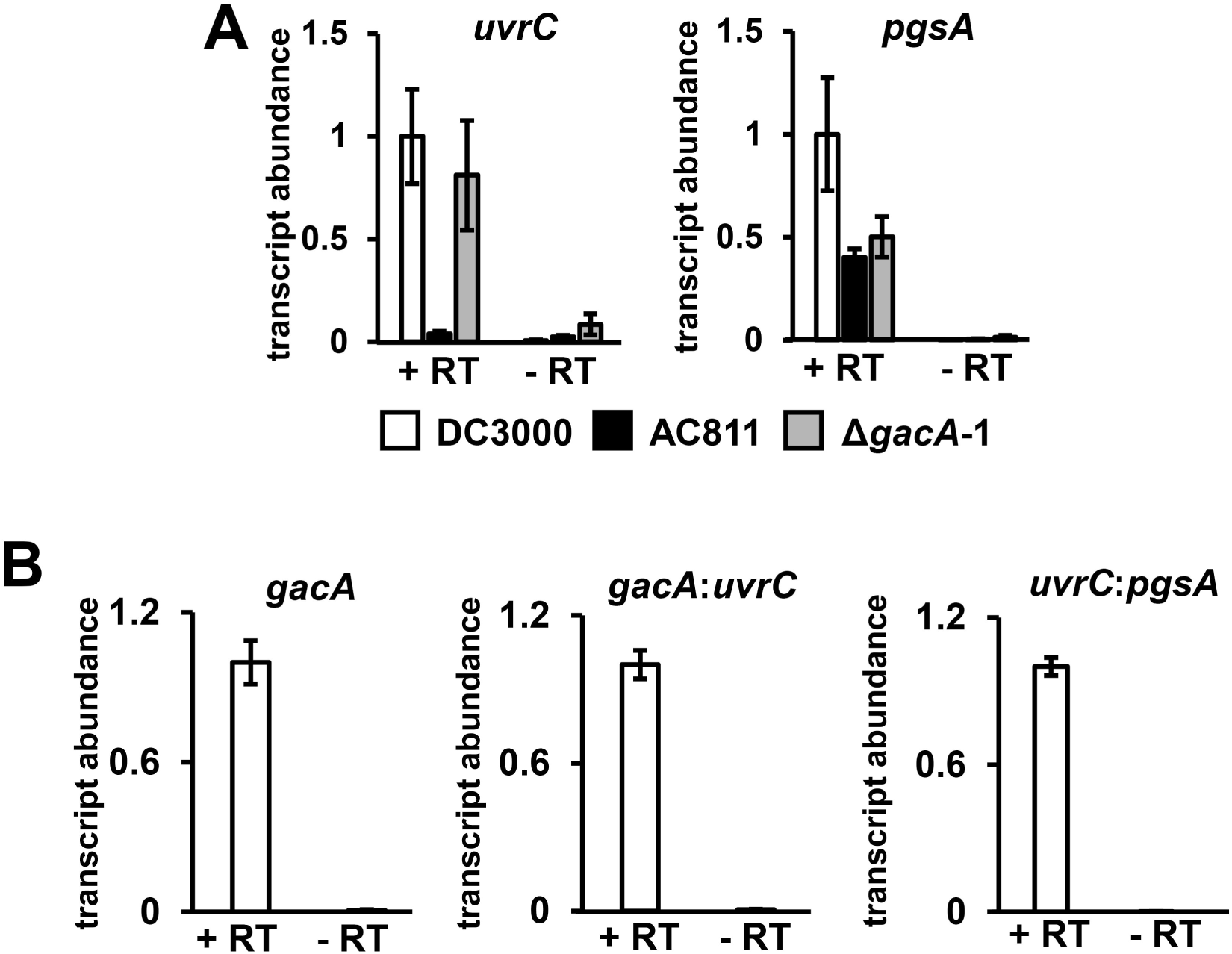
Reverse transcriptase-dependent quantification of *gacA* operon-associated transcripts. **(A)** RNA was extracted from DC3000, AC811, and Δ*gacA*-1, and cDNA synthesis performed with (+RT) or without reverse transcriptase (-RT). Shown is quantitative RT-PCR analysis of *gacA, uvrC*, and *pgsA* transcripts in these samples. Graphed are means ± SE of *uvrC* and *pgsA* transcript abundance normalized to *gyrA* from corresponding +RT samples and calculated relative to transcript levels measured in DC3000 (+RT). Data are pooled from two independent experiments with two technical replicates each; n = 4. Data for +RT samples are the same as shown in Fig. 2B. **(B)** Transcript abundance of *gacA, gacA:uvrC*, and *uvrC:pgsA* junctions from DC3000 cDNA synthesized with (+RT) or without (-RT) reverse transcriptase. Data are pooled from two independent experiments with two technical replicates each; n = 4.

**Fig S5.**
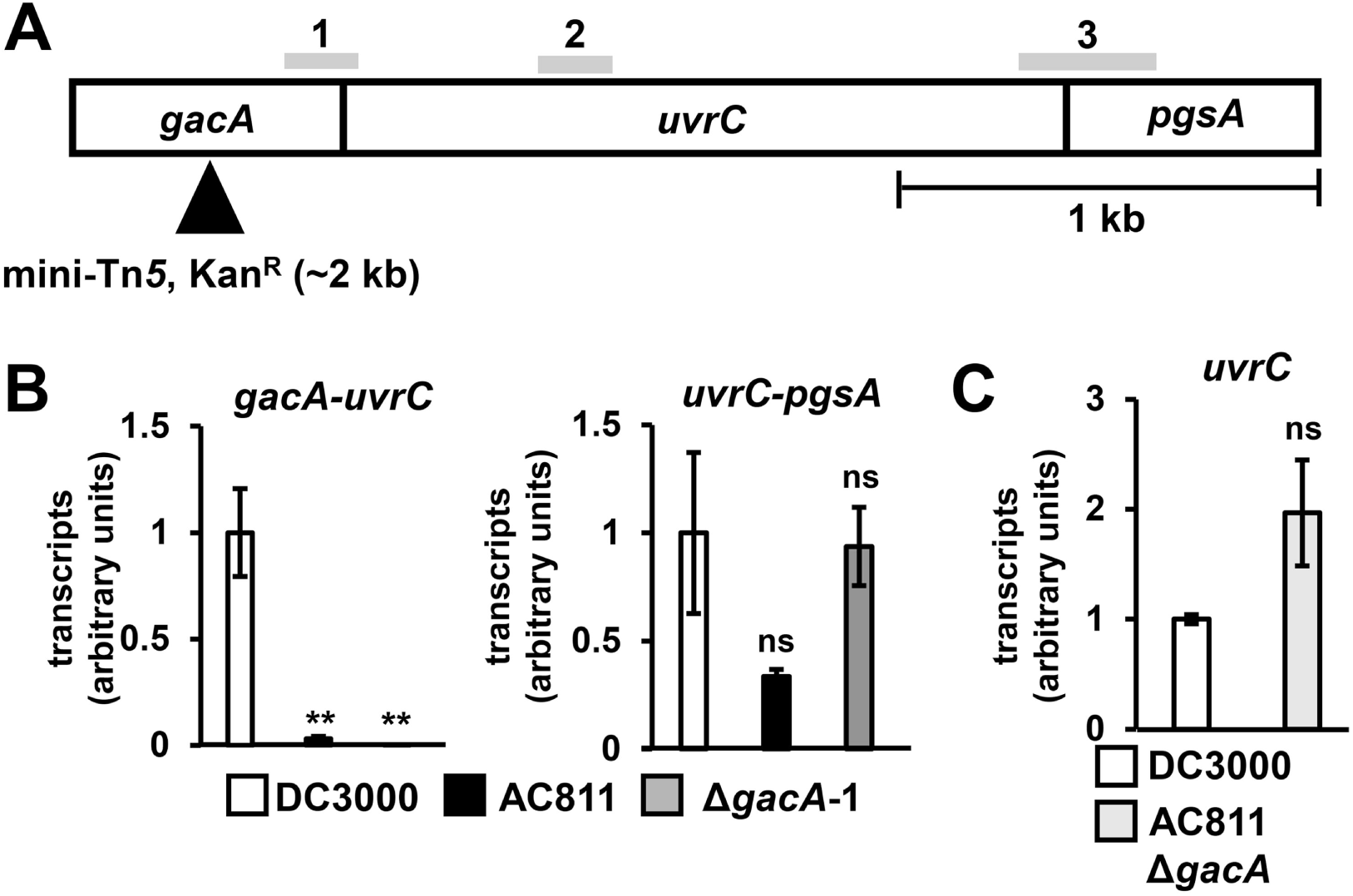
Polar effects of Tn*5* on *uvrC* and bicistronic *gacA-uvrC* and *uvrC-pgsA* transcript abundance. **(A)** Schematic of the predicted *gacA-uvrC-pgsA* operon in DC3000, with vertical bars indicating predicted translation start sites of *uvrC* and *pgsA*. Shading and numbering indicates regions targeted by qRT-PCR, with amplicons designated as follows: (1) *gacA:uvrC*; (2) *uvrC;* (3) *uvrC:pgsA*. **(B)** Abundance of *uvrC* transcripts was assessed by qRT-PCR using a *gyrA* reference gene as previously described. Values are normalized to DC3000. Graphed are means ± SE from data pooled across two independent experiments with two technical replicates each; n = 4. ns = no significant difference based on *t*-test (*p* > 0.05). **(C)** qRT-PCR analysis of transcripts containing *gacA-uvrC* or *uvrC-pgsA* junctions. Graphed are means ± SE from data pooled across two independent experiments with two technical replicates each; n = 4. \*\**p* < 0.01; ns = no significant difference (*p* > 0.05).

**Fig S6.**
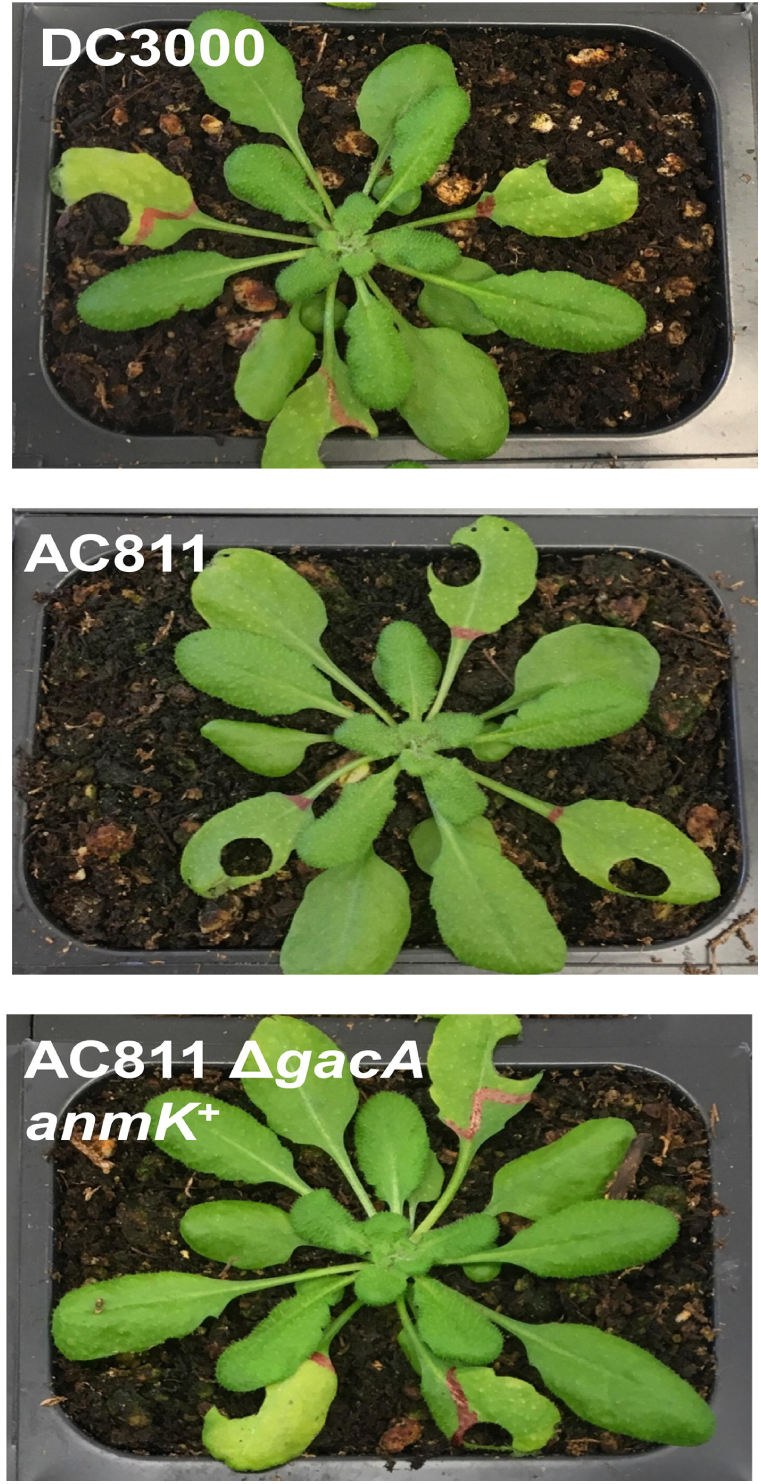
DC3000 and an AC811 Δ*gacA anmK*^+^ strain cause similar disease symptoms during infection of Arabidopsis. Arabidopsis leaves were syringe-infiltrated with DC3000, AC811, or AC811 Δ*gacA* carrying pME6010::*anmK*. Infected plants were photographed at 3 days post-infection (dpi). Circular punches on infected leaves are from sampling for bacterial cfu enumeration. Images are representative of symptoms observed in three independent experiments.

